# The Rho GTPase Rac1 mediates exercise training adaptations

**DOI:** 10.1101/2023.10.08.561442

**Authors:** Steffen H. Raun, Carlos Henriquez-Olguín, Emma Frank, Jonas Roland Knudsen, Mona S. Ali, Nicoline R. Andersen, Lisbeth L. V. Møller, Jonathan Davey, Hongwei Qian, Ana Coelho, Christian S. Carl, Christian T. Voldstedlund, Bente Kiens, Rikard Holmdahl, Paul Gregorevic, Thomas E. Jensen, Erik A. Richter, Lykke Sylow

## Abstract

Exercise training elicits tremendous health benefits; however, the molecular underpinnings are poorly understood. As one of the most regulated groups of proteins following acute exercise in human muscle, Rho GTPases are unexplored candidates for mediating the beneficial effects of exercise. The Rho GTPase Rac1 was activated during multiple exercise modalities and remained elevated hours after resistance exercise in human muscle. Inducible muscle-specific Rac1 knockout (Rac1 imKO) mice, displayed attenuated muscle protein synthesis, glycogen resynthesis and p38 MAPK signaling in recovery from contractions. Exercise training upregulated Rac1 protein content in human and mouse muscle. Overexpression of hyperactive Rac1 elevated reactive oxidant species production during exercise yet did not induce a trained muscle phenotype. In Rac1 imKO mice, the improvements in running capacity and muscle mass after exercise training were diminished. Using gain- and loss-of-function mouse models and human muscle biopsies, we identify Rac1 as a regulator of exercise training adaptions.

**Highlights:**

- Various exercise modalities activate Rac1 signaling in human skeletal muscle.
- HSP27, MNK1, and CREB are Rac1-dependent contraction-responsive targets in muscle.
- Post-contraction protein synthesis requires Rac1 but not NOX2.
- Rac1-NOX2 signaling is necessary for post-contraction glycogen resynthesis.
- Exercise training increases Rac1 protein content in human and mouse muscles.
- Rac1 mediates critical adaptations to exercise training.

## Introduction

Regular exercise training leads to multiple health benefits and can prevent and/or treat lifestyle- and age-related diseases^1^. The beneficial effect of exercise is mediated through signaling pathways that result in molecular adaptations, particularly within skeletal muscle, leading to increased muscle mass and function^2–4^. Targeting such mechanisms could potentially improve health^5^. However, the advancement of this potential is hampered by the lack of a comprehensive understanding of the mechanisms that improve muscle health in response to exercise.

Enrichment pathway analyses of the phospho-proteome of skeletal muscle indicate that Rho GTPases are one of the major groups of proteins regulated in the recovery after intense exercise^6^. The Rho family GTPase Rac1 (Ras-related C3 botulinum toxin substrate 1) is an exciting candidate for the regulation of muscle mass and function. Rac1 is known for its critical cellular importance in actin cytoskeleton regulation^7^ and the activation of the NADPH oxidase 2 complex (NOX2)^8,9^. In striated muscle, this GTPase is also necessary for cardiac hypertrophy^10–12^ and controls the metabolic response during acute exercise in mouse skeletal muscle^13–15^. Accordingly, Rac1 is activated by exercise in mouse and human skeletal muscle^16^. Here, Rac1 controls intramyocellular glucose uptake^15^ and NOX2-dependent reactive oxygen species (ROS) production^13,14^. Thus, there is growing evidence supporting the role of Rac1 during acute exercise, however it is unknown whether Rac1 is involved in the regulation of intramyocellular signaling in the recovery following exercise and the long-term adaptations to exercise training.

Consequently, we hypothesized that skeletal muscle Rac1 regulates post-exercise signaling to mediate the beneficial health effects of training in muscle. Using gain- and loss-of-function mouse models in combination with human muscle biopsies, we find that Rac1 regulates post-exercise intramyocellular signaling via p38MAPK-HSP27-CREB. Moreover, Rac1 was necessary for protein synthesis and glycogen resynthesis in recovery from muscle contraction. Ultimately, skeletal muscle Rac1 was required for critical adaptations to exercise training.

## Results

### Various exercise modalities activate Rac1 in human skeletal muscle

To obtain a temporal and modality-specific understanding of exercise-induced Rac1 activation, we determined Rac1 signaling in human skeletal muscle before, during, and in the recovery of three different exercise modalities^6^: endurance, sprint, and resistance exercise (Figure 1A). Group I PAKs (PAK1-3) bind active GTP-bound Rac1^17^. Active Rac1-binding to PAK1 and PAK2 elicits a conformational change in the kinases allowing for autophosphorylation of PAK1 on T423. This phosphorylation relieves PAK1 autoinhibition^18^ and phosphorylated (p)PAK1 T423 therefore serves as an indirect readout of Rac1 activity as used previously^19–21^.

**Figure 1:**
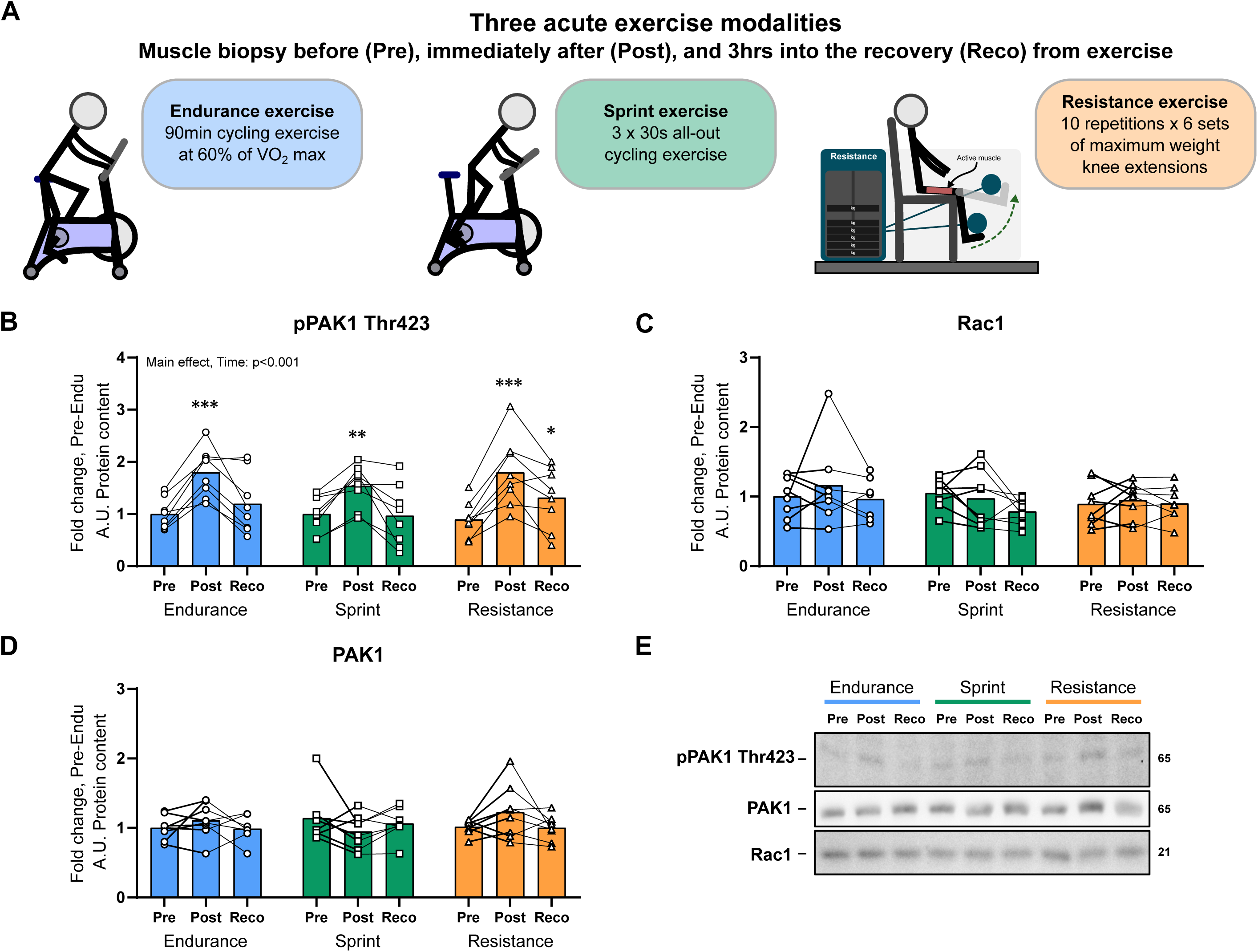
Multiple exercise modalities activate Rac1 signaling in human muscle. (A) Rac1 signaling in skeletal muscle (vastus lateralis) of young healthy men before (Pre), immediately after (Post), and 3 hours into the recovery (Reco) from endurance, sprint, or resistance exercise: (B) pPAK1 T423, (C) Rac1, and (D) PAK1. Representative blots are shown in (E). n= 8. Significant differences from pre are indicated; */**/*** = p<0.05/ p<0.01/ p<0.001. Data are presented as mean + SEM incl. individual values. Samples from the same individual are connected with lines.

Immediately following exercise cessation, pPAK1 T423 was elevated 50-80% independent of exercise modality (Figure 1B). These results expand upon our previous findings that Rac1 is activated in human skeletal muscle in response to uphill walking^16^. In the recovery phase, pPAK1 T423 remained 30% elevated 3 hours after resistance exercise, while the phosphorylation returned to pre-exercise levels in endurance and sprint exercise (Figure 1B). Expectedly, total Rac1 and PAK1 protein content were not changed after exercise or in recovery (Figure 1C-D). Representative blots are shown in Figure 1E. These findings show that endurance, sprint, and resistance exercise acutely activate Rac1 signaling in human skeletal muscle, which remains activated in the recovery phase following resistance exercise.

### HSP27, MNK1, and CREB are novel Rac1-dependent contraction-responsive targets in muscle

Having found that Rac1 was activated in response to various exercise modalities and remained elevated in the hours after resistance exercise in human skeletal muscle, we next sought to determine the intramyocellular signaling events activated by Rac1 in response to muscle contraction. To this end, we used an *in-situ* contraction protocol in mice which is known to increase intramyocellular signaling toward muscle protein synthesis in the recovery phase^22^. This protocol robustly activated Rac1 signaling inferred from a 45% increase in pPAK1 T423 (Figure 2A). Acute contraction-induced Rac1 activation occurred independent of mTORC1 as pPAK1 T423 was unaffected by rapamycin treatment (Figure 2A). Next, we used inducible skeletal muscle-specific Rac1 knockout (Rac1 imKO) and control littermate mice ^15,16^ with the objective to identify Rac1-dependent contraction-induced signaling mechanisms in recovery following muscle contraction (Figure 2B, Suppl. Figure 1A).

**Figure 2:**
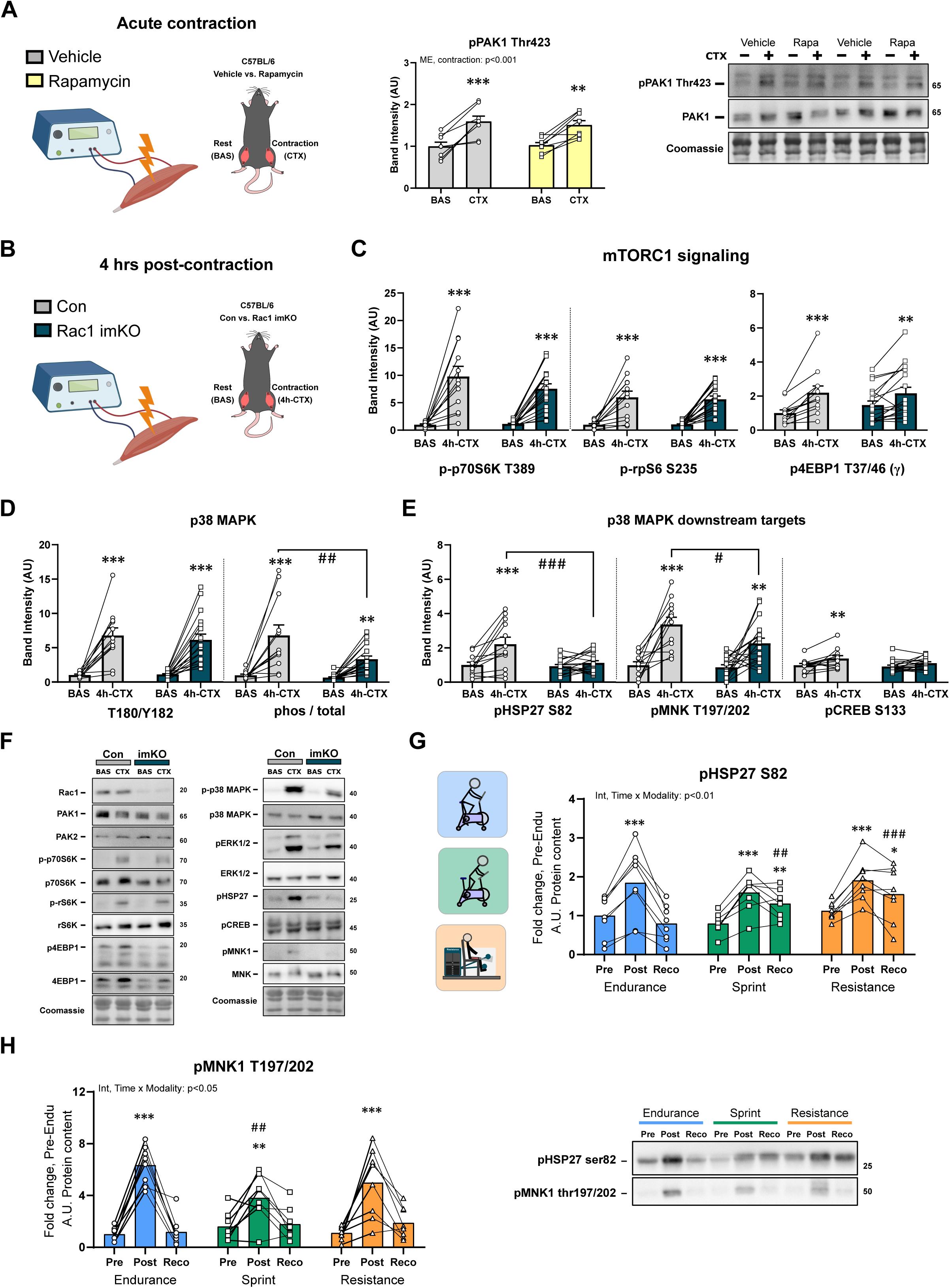
HSP27, MNK1, and CREB are Rac1-dependent contraction-responsive targets in muscle. (A) pPAK T423 mouse quadriceps muscle in response to *in situ* contraction of with or without rapamycin treatment. (B) Experimental setup of the investigation of post-contraction (4 hours, 4h-CTX) intramyocellular signaling in control (Con) and inducible muscle-specific Rac1 knockout (Rac1 imKO) mice. The effect of *in situ* contraction on: (C) mTORC1 signaling, (D) p38 MAPK phosphorylation, and (E) p38 MAPK signaling. Representative blots (F). (G-H) The effect of endurance, sprint, or resistance exercise in human skeletal muscle on pHSP S82 and pMNK1 T197/202. Acute contraction in mice (A): n=8. Post-contraction in mice (B-G), Control mice: n=12, Rac1 imKO mice: n=18. Humans (H): n=8. Significant differences basal leg vs. contraction are indicated; */**/*** = p<0.05/ p<0.01/ p<0.001. Significant differences Rac1 imKO vs. control are indicated as; #/# # /# # # = p<0.05/p<0.01/p<0.001. For human data, significant differences pre vs. Post/Reco are indicated; */**/*** = p<0.05/ p<0.01/ p<0.001, while differences between exercise modalities within time-point are indicated by; # #/# # # = p<0.01/p<0.001. Data are presented as mean+SEM incl. individual values.

As expected, and previously reported^22,23^, canonical mTORC1 signaling towards p70S6K (T389)/rS6 (S235-S236) and 4EBP1 (T37-46) (Figure 2C) was increased 4 hours following muscle contraction (Figure 2C). The contraction-induced phosphorylation of the mTORC1 downstream targets was unaffected by Rac1 imKO (Figure 2C), suggesting that Rac1 is dispensable for the activation of mTORC1 in the recovery from muscle contraction. Mitogen-activated protein kinases (MAPKs) are another group of proteins that have been implicated in contraction-induced muscle signaling^24^. The phosphorylation of p-p38MAPK T180/Y182 robustly increased 4 hours into recovery from *in situ* contraction irrespective of genotype (Figure 2D). Yet, we noted that Rac1 mKO mice had 50% increased total p38MAPK protein content (Suppl. Figure 1B). When related to the total p38 MAPK protein content, Rac1 mKO mouse muscle exhibited 50% reduced p-p38 MAPK T180/Y182 compared to control littermates (Figure 2D). This difference was specific to p38 MAPK, as ERK1/2 MAPK phosphorylation was not affected by Rac1 imKO (Suppl. Figure 1B). Total protein contents are presented in Suppl. Figure 1B.

To better understand the downstream consequences of lower relative contraction-stimulated p-p38 MAPK T180/Y182 in Rac1 imKO muscle, we next determined the phosphorylation of suggested p38MAPK targets. These include HSP27 (Heat-shock protein 27, also known as HSPB1)^25^, MNK1 (MAP kinase-interacting serine/threonine-protein kinase 1)^26,27^, and CREB (cAMP response element-binding protein)^28^. These proteins are known to be involved in vital cellular processes including gene transcription, protein translation, and functioning as chaperones. Contraction increased the phosphorylation of all these suggested p38MAPK targets in recovery in control mice (Figure 2E). In Rac1 imKO muscle, pHSP27 S82 was abolished in recovery from contraction (Figure 2E). In addition, both pMNK1 T197/202 and pCREB S133 were lower in Rac1 imKO compared to controls in recovery from contraction (Figure 2E). Representative blots are shown in Figure 2F. The expression of genes involved in metabolism and muscle growth were increased in recovery independent of Rac1 (Suppl. Figure 1C). Intrigued by the findings that HSP27, MNK1, and CREB are novel Rac1-dependent contraction-responsive targets in muscle, we turned back to verify the regulation of these signaling molecules in human skeletal muscle in response to endurance, sprint, and resistance exercise. In alignment with the mouse data, pHSP27 S82 (Figure 2G), pMNK1 T197/202(Figure 2H), and pCREB S133 (published in^29^) were increased acutely in response to exercise in all three exercise modalities. In addition, pHSP27 S82 was elevated 3 hours into the recovery of sprint and resistance exercise (Figure 2G). In combination with the data obtained from Rac1 imKO mice, these findings suggest that HSP27, MNK1, and CREB Rac1-dependent targets in the recovery from contraction in mouse and human skeletal muscle.

### Post-contraction protein synthesis and glycogen replenishment require Rac1

The rate of muscle protein synthesis increases significantly following a single bout of resistance exercise in humans^30^, which after repeated sessions ultimately results in muscle hypertrophy. This response is partially mediated by mTORC1, however, other signaling events may contribute^23,31–33^. Because of Rac1’s requirement for cardiac hypertrophy^10–12^, we hypothesized that Rac1 would be involved in the increase in skeletal muscle protein synthesis in the recovery from exercise. We, therefore, measured puromycin incorporation in skeletal muscle four hours after muscle contraction in Rac1 imKO and littermate control mice (Figure 2B). In line with our hypothesis and Rac1’s role in the heart^10–12^, Rac1 imKO mice showed decreased protein synthesis four hours after contraction compared to control mice (Figure 3A, Suppl. Figure 1D). Another anabolic event occurring in the recovery from exercise, is the replenishment of glycogen stores^34^. Rac1 imKO muscles failed to increase muscle glycogen in the recovery, where contraction led to 40% increased muscle glycogen content four hours after contraction in the control mice (Figure 3B). Collectively, the findings indicate an essential role of Rac1 in protein synthesis and glycogen resynthesis in recovery from muscle contraction.

**Figure 3:**
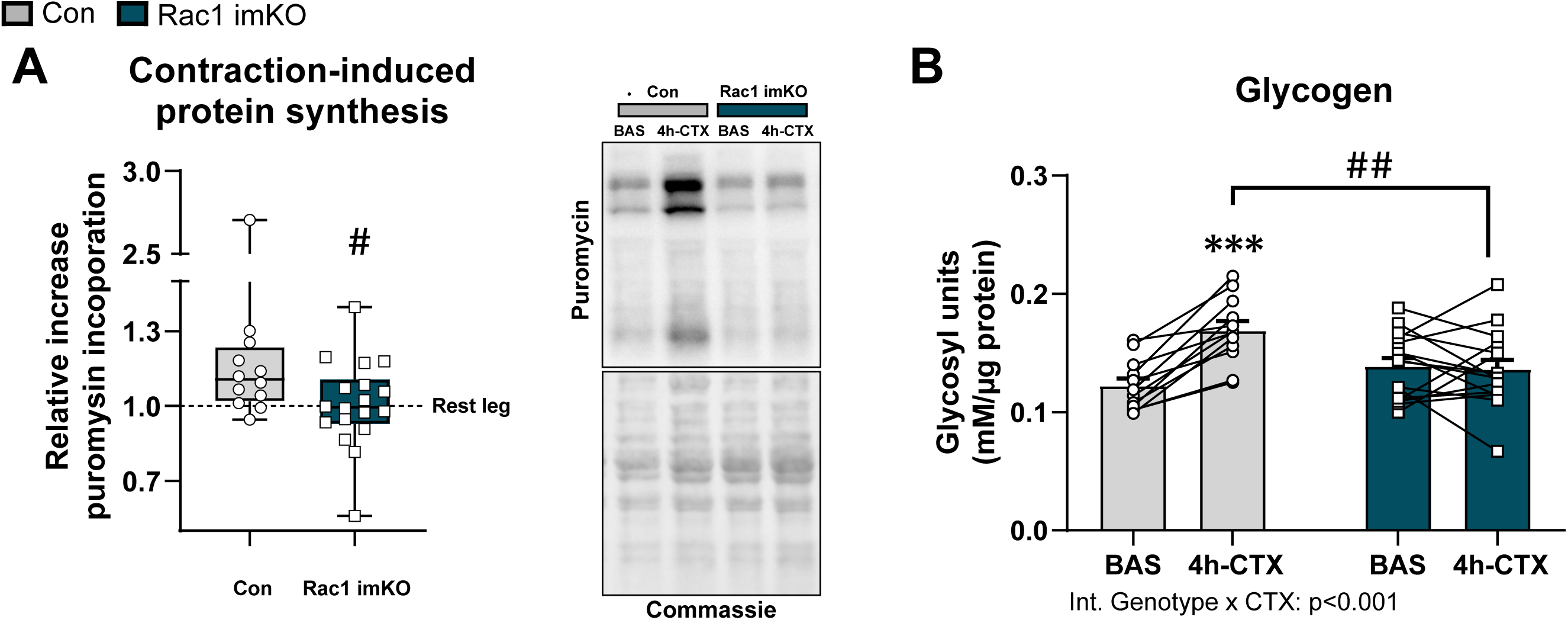
Post-contraction protein synthesis and glycogen resynthesis require Rac1. (A) Relative increase in protein synthesis (puromycin incorporation measured by western blot) and (B) glycogen synthesis 4 hours after contraction (quadriceps muscle) in control (Con) and inducible muscle-specific Rac1 knockout mice (Rac1 imKO). Control mice: n=12, Rac1 imKO mice: n=16-18. Significant differences Rac1 mKO vs. Con are indicated as; # = p<0.05, # # # = p<0.001. Data are presented as boxplots or bar plots and individual values.

### Rac1 affects post-contraction signaling and protein synthesis independent of NOX2 activity

Rac1 is necessary for the assembly and activation of the NOX2 complex to produce ROS during acute exercise^13^. This mechanism is required for Angiotensin-II induced cardiac hypertrophy ^35^, yet it is unexplored in skeletal muscle contraction-induced hypertrophic signals. Additionally, Rac1 is also necessary for exercise-stimulated glucose uptake in muscle via NOX2- derived ROS^13^.

To test whether Rac1 is involved in post-contraction p38 MAPK-signaling, protein synthesis, and glycogen resynthesis via its involvement in NOX2 assembly and ROS production, we used a mouse model carrying a loss-of-function mutation in the *Ncf1* gene. This mutation leads to an impaired function of the regulatory NOX2 subunit, p47^phox 36,37^ (Figure 4A) (*Ncf1\** mice). Similar to Rac1 imKO mice, in *Ncf1\** mouse muscle, NOX2-dependent production of ROS is blocked during acute exercise^13^. The *Ncf1\** mouse is thus useful for determining the role of Rac1- dependent NOX2-induced ROS production in the regulation of post-contraction signaling events. We found that protein synthesis (puromycin incorporation) was equally increased 4 hours into recovery from contraction in *Ncf1\** mice compared to control mice (Figure 4B). Interestingly, the increase in muscle glycogen in recovery was abolished in muscle lacking NOX2 activity (Figure 4C). In relation to signaling event, mTORC1 signaling, p38 MAPK-signaling, and downstream targets of p38 MAPK all increased their phosphorylation to the same extent in both genotypes (Figure 4D). Representative blots are shown in Figure 4E.

**Figure 4:**
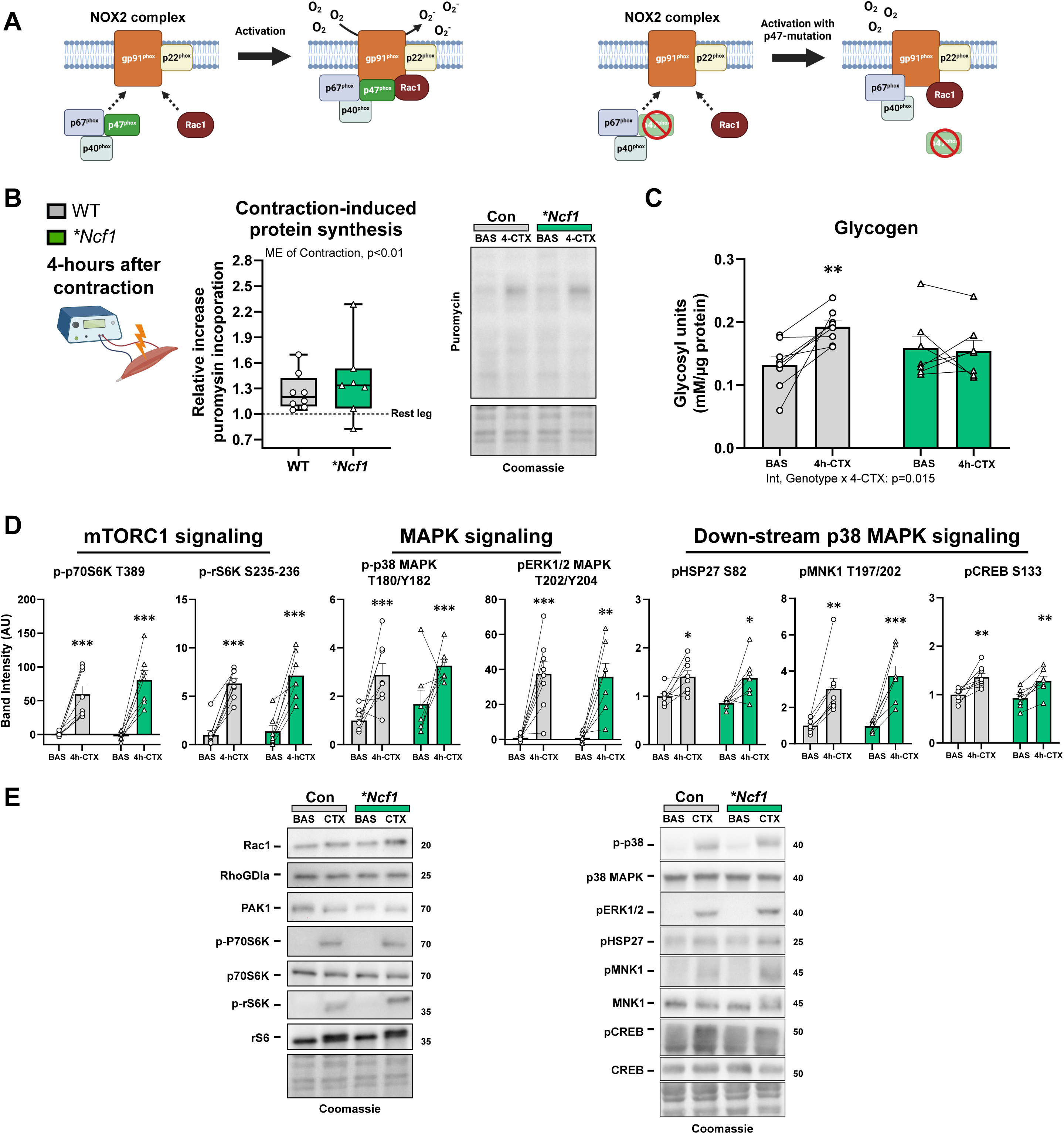
Rac1 affects post-contraction protein synthesis independent of NOX2 activity. (A) Illustration of the activation of the NOX2 complex and subsequent O2^-^-production with and without p47phox mutation (*\*Ncf1*). (B) Relative increase in protein synthesis (puromycin incorporation measured by western blot) in quadriceps muscle from control mice and mice carrying a mutation in the *\*Ncf1* gene (p47^phox^). (C) Muscle glycogen after contraction. (D) Phosphorylation of mTORC1 substrates, p38 MAPK and ERK1/2 MAPK, and p38 MAPK substrates. (E) Representative blots. Control: n=8, *\*Ncf1*: n=7. Significant differences from basal vs. contraction are indicated; */**/*** = p<0.05/ p<0.01/ p<0.001. Data are presented as boxplots or mean+SEM incl. individual values.

Taken together these data suggest that Rac1 mediates p38 MAPK signaling and protein synthesis in recovery independent of NOX2-mediated ROS production. In contrast, Rac1 regulates post-contraction muscle glycogen resynthesis via the NOX2 complex, a signaling pathway shown to be involved in acute exercise-stimulated glucose uptake ^13,15^.

### Exercise training elevates Rac1 protein content in human and mouse skeletal muscle

Having established Rac1 as a critical regulator of contraction-induced signaling and adaptations into recovery, we next wanted to know if Rac1 protein content was regulated by long-term exercise training. Accordingly, we analyzed muscle biopsy samples from lean healthy men before and after a single-leg knee-extensor exercise training intervention (published in^38^). Three weeks of exercise training led to a 30% increase in Rac1 protein content, which was unaltered in the non-trained control leg (Figure 5A). Similarly, Rac1 protein content was also modest increased in mouse gastrocnemius muscle following voluntary wheel running exercise training (Figure 5B). Thus, Rac1 is a novel exercise training responsive protein in human and mouse skeletal muscle that could potentially mediate exercise training adaptations in skeletal muscle.

**Figure 5:**
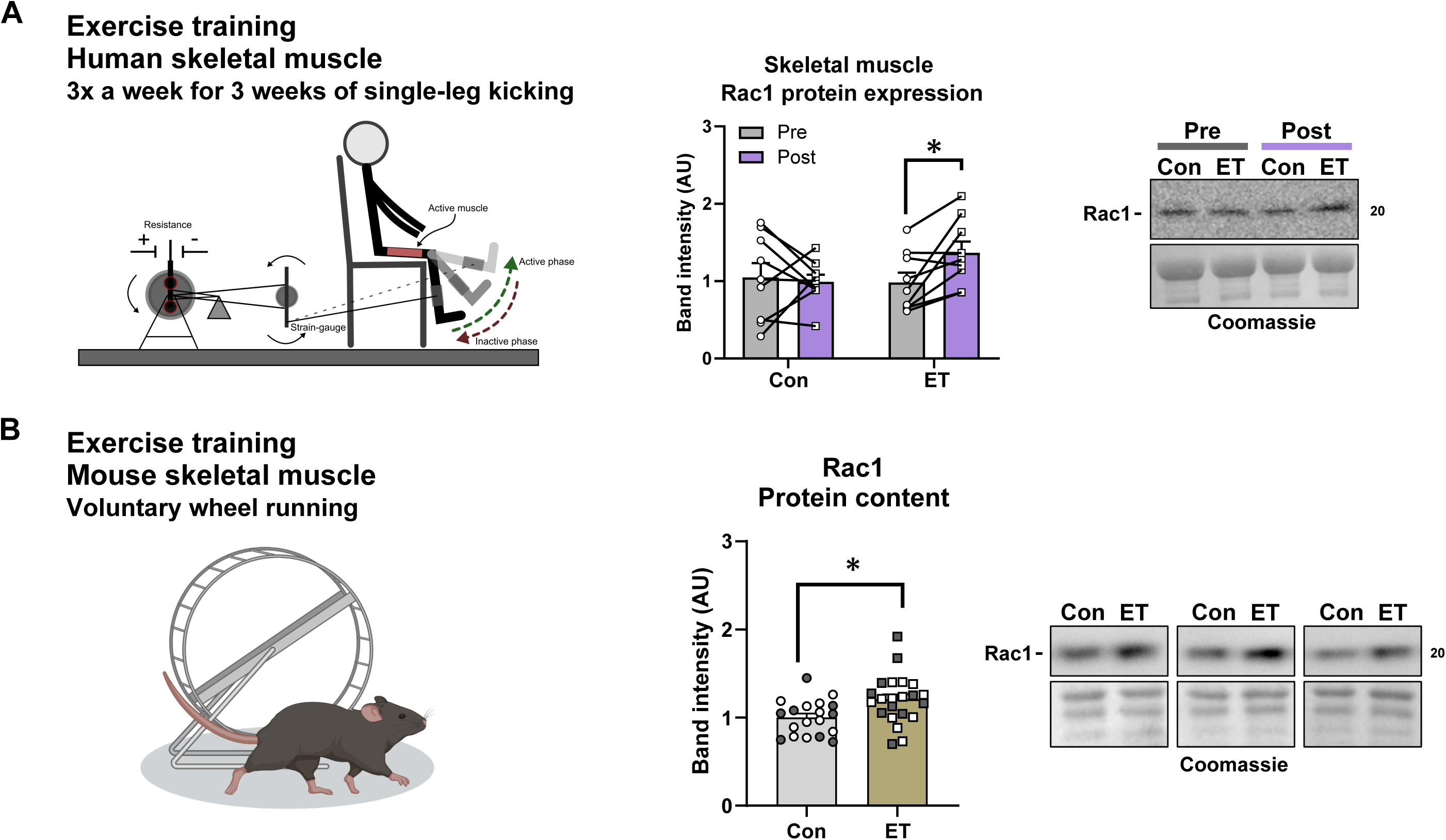
Exercise training increases skeletal muscle Rac1 protein content. (A) Effect of single-leg exercise training (ET) on Rac1 protein content in human skeletal muscle (vastus lateralis). (B) Effect of voluntary wheel running ET on Rac1 protein content in mouse gastrocnemius muscle (Dark dots=12 weeks intervention, clear dots=6 weeks intervention). Human study: n=9. Mouse study: Control: n=19, ET: n=22. Significant of ET; * = p<0.05. Data are presented as mean+SEM incl. individual values, connected when applicable (pre vs. post).

### Rac1 mediates critical adaptations to exercise training

To determine the role of skeletal muscle Rac1 in exercise training adaptations, we conducted a 12 weeks voluntary wheel running exercise training intervention, which induces exercise training adaptation in skeletal muscle of mice^39–42^ (Figure 6A). Prior to the intervention, untrained Rac1 imKO mice had similar exercise capacity on a treadmill compared to littermate controls (Suppl. Figure 2A). While blood glucose increased similarly during a treadmill running performance test in both genotypes, blood lactate increased 30% only in control mice, suggesting a lower contribution of glycolysis in the Rac1 imKO mice during exercise (Suppl. Figure 2B). When challenged to an endurance treadmill exercise test, untrained Rac1 imKO mice showed a 25% reduction in running time compared to control mice (Suppl. Figure 2C).

**Figure 6:**
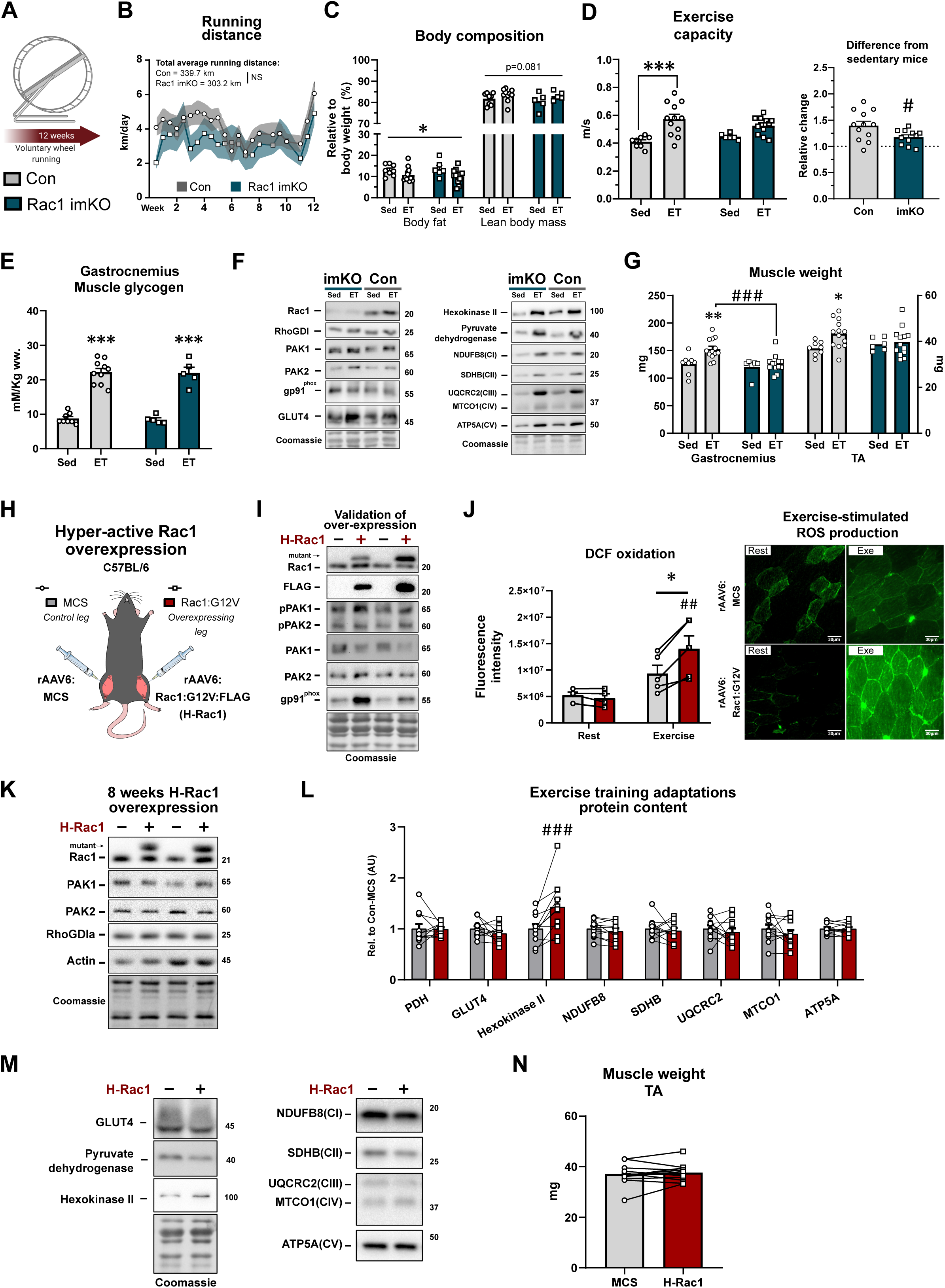
Rac1 mediates critical adaptations to exercise training. (A) Lack of skeletal muscle Rac1 during exercise training was investigated using inducible skeletal muscle-specific Rac1 knockout mice (Rac1 imKO) compared to control littermate mice (Con). (B) Running distance during the intervention. Effect of ET on (C) body composition, (D) exercise capacity, (E) muscle glycogen (gastrocnemius), (F) protein content measured by western blotting in gastrocnemius muscle, and (G) muscle weight. (H) hyperactive Rac1 (H-Rac1) was overexpressed using recombinant adeno-associated virus. (I) Immunoblotting validation of H- Rac1 overexpression. (J) DCF-oxidation (oxidant production) at rest and during exercise in tibialis anterior muscle. (K) Effect of 8 weeks of H-Rac1 overexpression. (L-M) Effect of H-Rac1 on muscle protein content measured by immunoblotting in gastrocnemius muscle. (N) Effect of H-Rac1 on muscle weight. For control mice: n=8-13. For Rac1 imKO mice: n= 5-13. For H-Rac1 studies: n=4-10. Significant differences in acute exercise / ET vs. SED control are indicated as; * = p<0.05, ** = p<0.01, *** = p<0.001. Significant differences Rac1 imKO or H-Rac1 overexpression vs. respective control are indicated as; # # = p<0.01, # # # = p<0.001. NS= non-significant. Data are presented as mean + SEM incl. individual values, paired samples connected with lines when applicable.

Rac1 imKO and control littermate mice ran similar daily distances (Figure 6B), and exercise training similarly lowered body fat by 20% in both genotypes (Figure 6C). The exercise training intervention expectedly ^39,41^ increased exercise capacity by 45% in control mice. However, the Rac1 imKO mice did not improve their exercise capacity compared to untrained Rac1 imKO mice (Figure 6D). Thus, the change in exercise capacity compared to non-trained mice was significantly reduced in Rac1 imKO compared to control mice (Figure 6D). This was observed despite similar exercise training-induced elevation (155%) of muscle glycogen between genotype (Figure 6E).

Expectedly, Rac1 protein content was reduced by 80% in Rac1 imKO compared to control mice with no effect on cardiac muscle Rac1 protein content (Figure 6F, bar plots are presented in Suppl. Figure 3A). To determine the exercise training adaptations in skeletal muscle, we next measured the protein content of proteins involved in Rac1 signaling and muscle metabolism. In gastrocnemius muscle, the kinases down-stream of Rac1, PAK1 and PAK2, were both increased in Rac1 imKO muscles, where PAK2 muscle protein content increased (50%) with exercise training in control mice (Figure 6F, Suppl. Figure 3B). In contrast to the down-stream effectors, the Rac1 inhibitor, Rho guanine dissociation inhibitor α (RhoGDIα)^43^, was neither affected by Rac1 imKO nor exercise training (Figure 6F, Suppl. Figure 3B). GLUT4 protein content increased (+35%) in control mice, but not in Rac1 imKO muscles after exercise training (Figure 6F, Suppl. Figure 3B). Lastly, Pyruvate dehydrogenase E1a (PDH), Hexokinase II, and electron transport chain proteins were all increased (+50% to +185%) in response to exercise training independent of Rac1 (Figure 6F, Suppl. Figure 3B).

Another key feature of the beneficial effects of exercise is muscle hypertrophy^44,45^. Interestingly, lack of Rac1 in skeletal muscle diminished training-induced muscle growth compared to control mice, which increased muscle mass by 20% and 15% in the gastrocnemius and tibialis anterior, respectively (Figure 6G). This aligns with our results showing that Rac1 was required for protein synthesis in recovery from muscle contraction (Figure 3). The difference in muscle growth was not due to genotypic differences of total protein content (measured in gastrocnemius muscle) of mTOR, p70S6K, or raptor (part of mTORC1), however rictor (part of mTORC2) increased with exercise training in a Rac1-dependent manner (Suppl. Figure 3C).

Having established Rac1 as a critical regulator of exercise training-induced adaptations, we next investigated whether transgenic activation of Rac1 would suffice to induce an exercise-trained muscle phenotype. For this, we constructed a recombinant adeno-associated viral vector (rAAV6) with muscle tropism^46^ to overexpress a hyperactive Rac1 mutant (H-Rac1) with an N-terminal FLAG-tag (Rac1:G12V) (Figure 6H). Rac1:G12V is a naturally occurring mutation located at the P-loop (phosphate-binding structure) of Rac1 and impairs Rac1-dependent hydrolysis of GTP, making this Rac1 mutant constitutively active^47–49^. H-Rac1 increased pPAK1 T423 (+30%) and pPAK2 T402 (+35%) despite reduced total PAK1 protein content (Figure 6I, Suppl. Figure 3D). This elevation was also observed during treadmill running (Figure 6I, Suppl. Figure 3D). In line with Rac1’s crucial role in NOX2-dependent exercise-induced ROS production, active H-Rac1 led to a 50% increase in oxidant production (DCFH fluorescence) during exercise, but not at rest (Figure 6J). Moreover, the catalytic subunit of the NOX2 complex, gp91^phox^, was elevated (+170%) in H-Rac1 muscles (Figure 6I, Suppl. Figure 3E).

After validating that our mutation worked as intended, we next overexpressed H-Rac1 for 8 weeks to investigate whether constitutive activation of Rac1 would lead to a trained muscle phenotype (Figure 6K). While Hexokinase II protein content increased (+30%) in response to H-Rac1, most exercise training-responsive proteins investigated were not affected by H-Rac1 overexpression in gastrocnemius muscle (Figure 6L-M). Moreover, H-Rac1 did not induce muscle hypertrophy (Figure 6N). Collectively, while activation of Rac1 is not sufficient to induce exercise training adaptations, Rac1 is required for critical adaptations to exercise training in skeletal muscle, including regulation of exercise capacity and muscle mass.

## Discussion

Skeletal muscle is remarkably adaptable. The present study demonstrates that skeletal muscle Rac1 regulates contraction-induced intramyocellular signaling and is required for multiple exercise training adaptations (Figure 7). These results not only unravel new exercise-induced, Rac1-dependent signaling pathways, but also adds to our understanding of the mechanisms that improve muscle health in response to exercise. Rac1’s role in the adaptive response to exercise was suggested by five major findings. First, we demonstrated that various exercise modalities activate Rac1 signaling in human skeletal muscle. Second, we identify HSP27, MNK1, and CREB as Rac1-dependent contraction-responsive targets in skeletal muscle. The functional roles of Rac1 were indicated by our third finding that Rac1 was necessary for increased protein synthesis and glycogen resynthesis in skeletal muscle in recovery from contraction. Fourth, exercise training elevated Rac1 protein content in human and mouse muscle, and finally, Rac1 mediated critical adaptations to exercise training, namely improvements in exercise capacity and increases in muscle mass.

**Figure 7:**
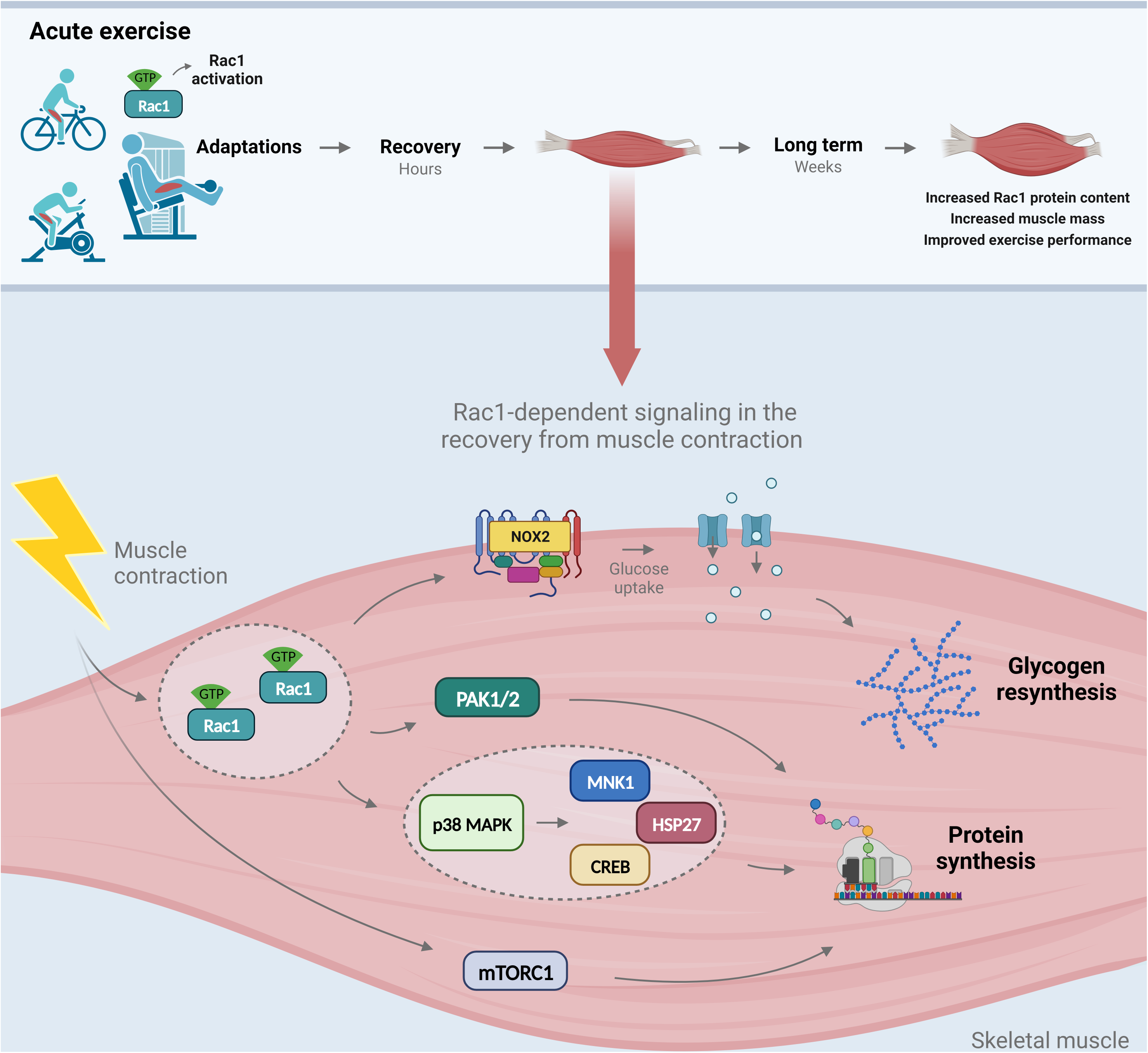
Graphical summary of key findings in current study.

We found that various exercise modalities, ranging from resistance to aerobic exercise training activated Rac1 in the vastus lateralis muscle of humans. These findings identify Rac1 as a canonical exercise-activated protein across different modalities and support previous research that uphill walking increased Rac1 GTP-binding and phosphorylation of Rac1’s target PAK^16^. The role of Rac1 in recovery is further supported by enrichment pathway analyses of the phospho-proteome in the recovery after intense exercise, which suggest that Rho GTPases are one of the most exercise-regulated groups of proteins^6^.

Exercise elicits multiple signals during and in recovery from exercise, ultimately leading to phenotypic changes in skeletal muscle. Our second major finding was that Rac1 played a key role in mediating these exercise-induced signals in recovery from contraction. Specifically, we identify the putative p38MAPK targets, HSP27, MNK1, and CREB as Rac1-dependent contraction-responsive targets in muscle. The mechanistic implications of Rac1 in muscle likely translate to humans, as we observed a robust increase in HSP27, MNK1, and CREB phosphorylation in all three exercise modalities. Moreover, in recovery in humans, HSP27 remained phosphorylated. Inconsistency between the phosphorylation of p38 MAPK and downstream targets in recovery between mouse and humans could be due to species differences, temporal differences, or differences in contraction protocols.

While it is currently unknown how Rac1 regulates p38 MAPK activity in the recovery from muscle contraction, it may involve Rac1’s implication in stretch-induced mechano-transduction during contraction^50^. Mechanistically, Rac1-induced signaling was independent of the key regulator of muscle size, mTORC1^51^, as this signaling pathway was not affected by the loss of Rac1 in muscle. Thus, while Rac1 has been shown to bind and regulate mTORC1 in non-muscle cells^52^, this does not seem to translate to mature skeletal muscle in response to contraction. Moreover, although NOX2 has previously been suggested to regulate p38 MAPK phosphorylation during acute treadmill running^13^, Rac1-mediated signaling in recovery from contraction was not affected in *Ncf1\** mouse muscle. Clues pertaining to the downstream effectors of Rac1-p38MAPK-associated signaling are in the existing body of literature. P38MAPK and CREB have previously been shown to induce *Pgc1α* gene expression following muscle contraction^53,54^ and p38MAPK can stimulate upstream transcription factors of *Pgc1α*, such as ATF2 and MEF2^55^. However, we observed no effect of Rac1 imKO on contraction-induced *Pgc1α* gene expression or other mitochondrial-related genes, as well as no effect on mitochondrial biogenesis after exercise training. Thus, these data collectively hint toward PGC1*α*-independent effects of Rac1. Aligning with the lower contraction-induced protein synthesis and abrogated HSP27 and CREB phosphorylation in Rac1 imKO mice, global KO of HSP27, led to lower body mass and smaller muscle fibers^56^, while overexpression of the CREB coactivators induced increased muscle mass^57^. In contrast, MNK1’ role in skeletal muscle is undefined. In non-muscle cells, MNK1 is implicated in the control of translation initiation^58^. However, double MNK1/2 Knockout (MNK1/2 dKO) in fibroblast does not affect general protein synthesis^59^, and global MNK1/2 dKO in mice does not affect body weight^60^, although muscle mass was not assessed. Lastly, increased expression of Rac1’s downstream target, PAK1, prevented cancer-induced muscle mass loss in mice^61^. Moreover, double KO of PAK1 (whole-body) and PAK2 (muscle-specific) mice have reduced muscle mass^62^ and late-onset myopathy^63^. Together these findings indicate that in response to muscle contraction, Rac1 regulates multiple proteins and pathways critically implicated in muscle mass control.

Those findings align with our third major finding, which revealed a functional role for Rac1 in increasing protein synthesis in recovery from contraction. We interpret these combined results to indicate that contraction-induced activation of Rac1 stimulates protein synthesis via a previously undescribed cascade involving Rac1, p38 MAPK, and its downstream signaling to PAK, HSP27, MNK, and CREB. This signaling cascade is independent of mTORC1-signaling and NOX2 activity. Noteworthy, these data contrast recent findings in diet-induced obese *Ncf1\** mice, where global lack of NOX2 activity prevented the increase in muscle hypertrophy after exercise training^64^. Overall, our findings supplement a growing body of evidence of mTORC1- independent regulation of muscle mass^23^.

Our fourth major finding was that Rac1 protein content was elevated in human and mouse skeletal muscle following exercise training. Finally, skeletal muscle Rac1 was required for both muscle hypertrophy and improvements in exercise capacity in response to voluntary exercise training in mice. These results align well with the reduction in protein synthesis in the recovery from contraction of Rac1 imKO mice, which should be expected to impair muscle mass gains in response to repeated exercise training. Importantly, overexpression of hyperactive Rac1 was insufficient to induce skeletal muscle training adaptations, including hypertrophy. These results are distinct from the hypertrophy-inducing phenotype of constitutively active Rac1 in cardiac muscle^10–12^. The role of Rac1 in mediating muscle growth in response to other stimuli, such as resistance-type exercise^65^ or amino acids will be an interesting avenue for future research.

Another critical exercise adaptation that was mediated by Rac1, was the increase in running capacity. Downstream of Rac1, this effect could be mediated by PAK, as whole-body PAK1 KO mice did not improve running capacity following 6 weeks of treadmill exercise training^66^. However, a limitation of that study was that whole-body PAK1 KO mice display cardiac remodeling defects ^66^. The lack of improvement in exercise capacity may also be coupled with Rac1’s recruitment for activation of the NOX2 complex. Indeed, *Ncf1\** mice, lacking NOX2 activity and ROS production during exercise, do not increase their exercise capacity after high-intensity interval training^67^. The importance of ROS in aerobic training adaptations likely translates to humans, where greater whole-body oxidative stress elicited greater improvements in endurance capacity after a training intervention^68^. However, the *Ncf1\** mouse, like the PAK1 KO mouse, is a whole-body transgenic mouse, thus it is not clear whether the lack of improvements in running performance is directly caused by a lack of skeletal muscle adaptations or due to organismal and/or cardiac adaptations.

Taken together, exercise activates intramyocellular signaling that confer remarkable health benefits. By analyzing skeletal muscle samples and multiple exercise modalities in humans, combined with Rac1 loss-of-function and hyperactive transgenic mutants, our study position Rac1 at the center of exercise training adaptations. Specifically, Rac1 in muscle was necessary for protein synthesis and glycogen resynthesis in the recovery of contraction. Rac1’s role in exercise adaptations manifested as a lower exercise capacity and muscle mass in response to training in Rac1 imKO mice. These data were translated to human skeletal muscle, where Rac1 signaling was acutely upregulated in multiple modalities, while Rac1 protein content increased after exercise training. Thus, we identify Rac1 as a requirement for both functional and molecular adaptations to exercise training.

## Additional information

### Author contributions (CRediT)

**Conceptualization**: SHR, LS, EAR. **Methodology**: SHR and LS. **Formal analysis**: SHR. **Writing – original draft preparation**: SHR and LS. **Investigation:** SHR, LS, CHO, EF, JRK, AC, LLVM, MSA, NRA, JD, HQ, TEJ. **Resources:** LS, EAR, CSC, CTV, CF, RH, JFPW, BK, PG. **Writing – review and editing**: SHR, LS, EAR, CHO, EF, JRK, AC, LLVM, MSA, NRA, CSC, CTV, CF, RH, JFPW, BK, TEJ. **Visualization**: SHR. **Supervision**: LS and EAR. **Project administration**: SHR and LS. **Funding acquisition:** LS, EAR, SHR. All authors are guarantors of this work and take responsibility for the integrity of the data and the accuracy of the data analysis.

### Grants

SHR was supported by the Independent Research Fund Denmark (#2030-00007A) and the Lundbeck Foundation (R380-2021-1451). LS was supported by the Novo Nordisk Foundation (NNF16OC0023418 and NNF18OC0032082), Independent Research Fund Denmark (4004-00233B and 9039-00170B), and the Carlsberg Foundation (CF21-0369). EAR was supported by the Novo Nordisk Foundation (NNF17OC0027274 and NNF18OC0034072). TEJ was supported by a Lundbeck Ascending Investigator project (R313-2019-643). LLVM was supported by a PhD fellowship from The Lundbeck Foundation (R208-2015-3388) and a Postdoctoral fellowship from The Lundbeck Foundation (R322-2019-2688). CHO and JRK were supported by a research grant from the Danish Diabetes Academy, which is funded by the Novo Nordisk Foundation (CHO: NNF17SA0031406). RH and AC were supported from Vetenskapsrådet (2019-01209).

## Acknowledgements

We acknowledge the skilled technical assistance of Betina Bolmgren and Irene Bech Nielsen, who have been a tremendous help during specific analysis for this study (August Krogh Section for Molecular Physiology, Department of Nutrition, Exercise and Sports, University of Copenhagen, Denmark). We would like to acknowledge Christian Frøsig and Jørgen F.P. Wojtaszewski (August Krogh Section for Molecular Physiology, Department of Nutrition, Exercise and Sports, University of Copenhagen, Denmark) for allowing us to investigate Rac1 protein content in their previously collected human exercise training samples^38^. We also acknowledge Peter Schjerling, PhD, who helped with genotyping of the Rac1 imKO mice (Center for Heathy Aging, University of Copenhagen). We would also like to acknowledge Prof. Grahame Hardie (University of Dundee, UK) for providing us with the PDH antibody used in the current study.

## Data and materials availability

All data needed to evaluate the conclusions in the paper are present in the paper and/or the Supplementary Materials. Additional data related to this paper may be requested from the corresponding authors (LS/SHR) upon reasonable request.

## Supplementary figure legends

**Suppl. Figure 1:**
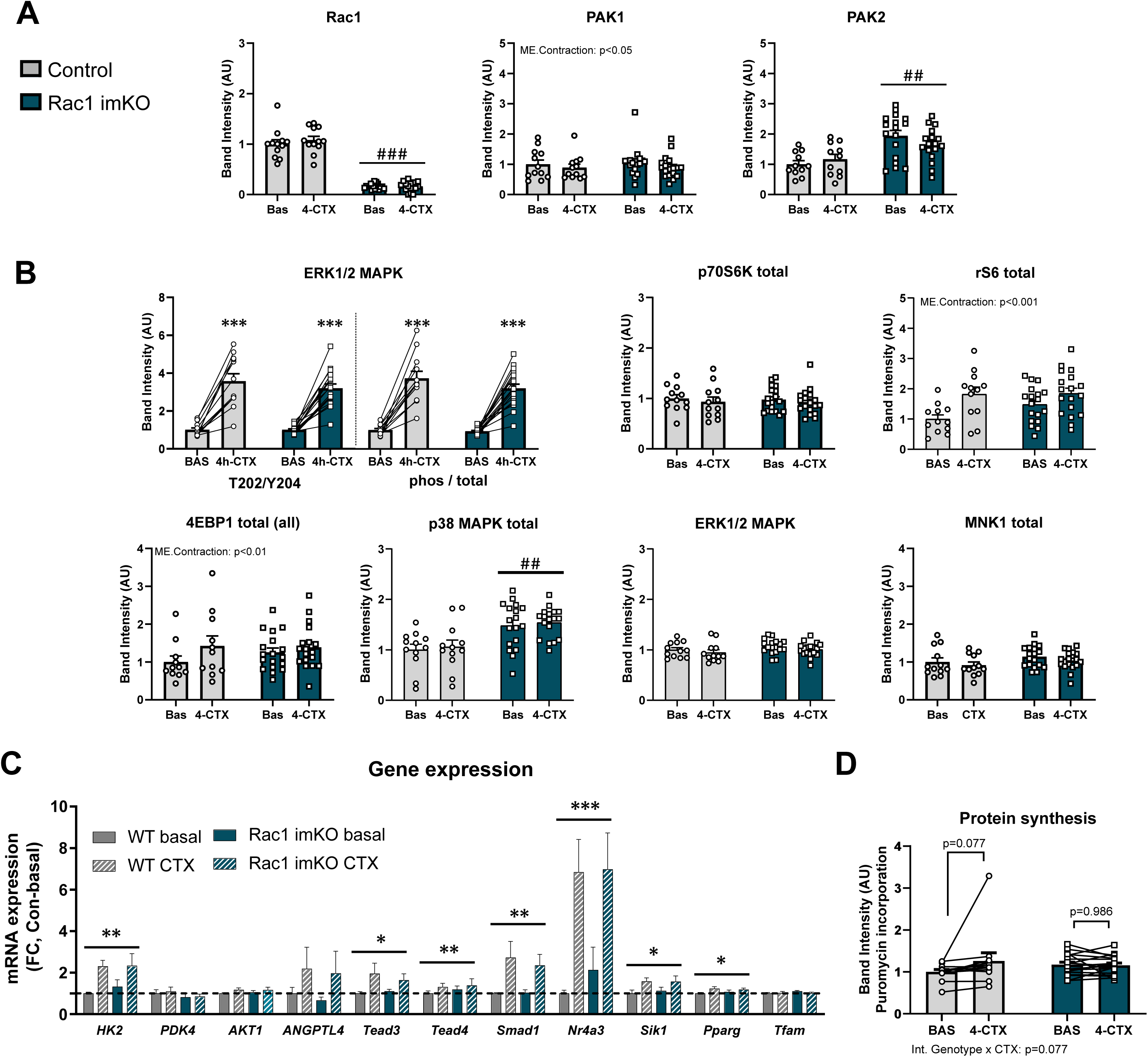
Data related to figure 2 and figure 3. Related to the investigation of post-contraction (4 hours, 4h-CTX) signaling events in control mice (Con) and inducible muscle-specific Rac1 knockout mice (Rac1 imKO). The effect of contraction on total protein content of: (A) Rac1, PAK1, PAK2, and (B) p-ERK1/2, p70S6K, rS6, 4EBP1, p38 MAPK, ERK1/2, MNK1. (C) Effect of contraction on gene expression of genes related to muscle growth and metabolism. (D) Raw data for protein synthesis in relation to Figure 3. Control mice: n=12, Rac1 imKO mice: n=18. Significant differences from basal leg vs. contraction are indicated; */**/*** = p<0.05/ p<0.01/ p<0.001. Significant differences of Rac1 imKO mice are indicated as; # # /# # # = p<0.01/p<0.001. Data are presented as mean+SEM incl. individual values.

**Suppl. Figure 2:**
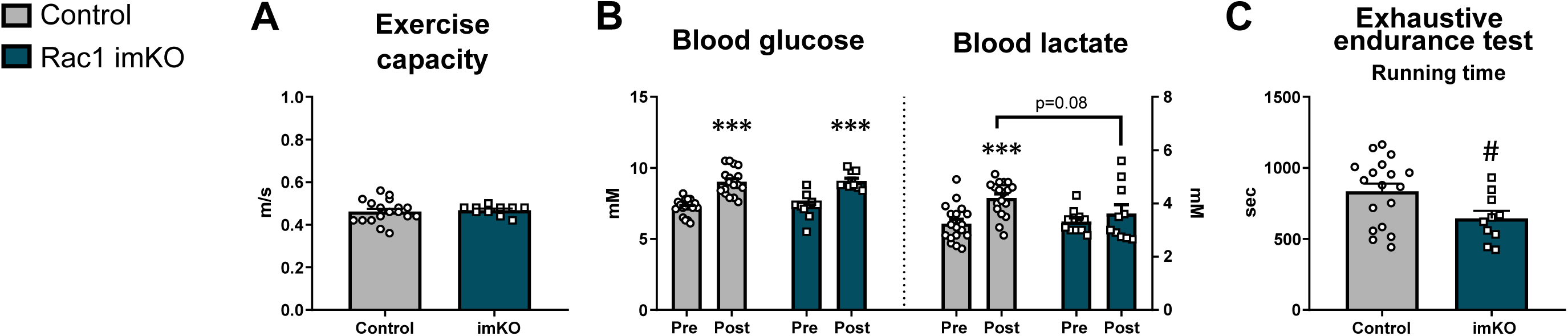
Data related to figure 6. (A) In untrained control mice and inducible muscle-specific Rac1 knock mice (Rac1 imKO), maximal running speed was measured during an incremental treadmill test, where blood glucose (B) and blood lactate (B) were measured before and after the test. The endurance capacity was measured during an treadmill running test (D). Control mice: n=18, Rac1 imKO mice: n=10. Significant differences of exercise are indicated as; *** = p<0.001. Significant differences of Rac1 imKO are indicated as; # = p<0.05. Data are presented as mean + SEM incl. individual values.

**Suppl. Figure 3:**
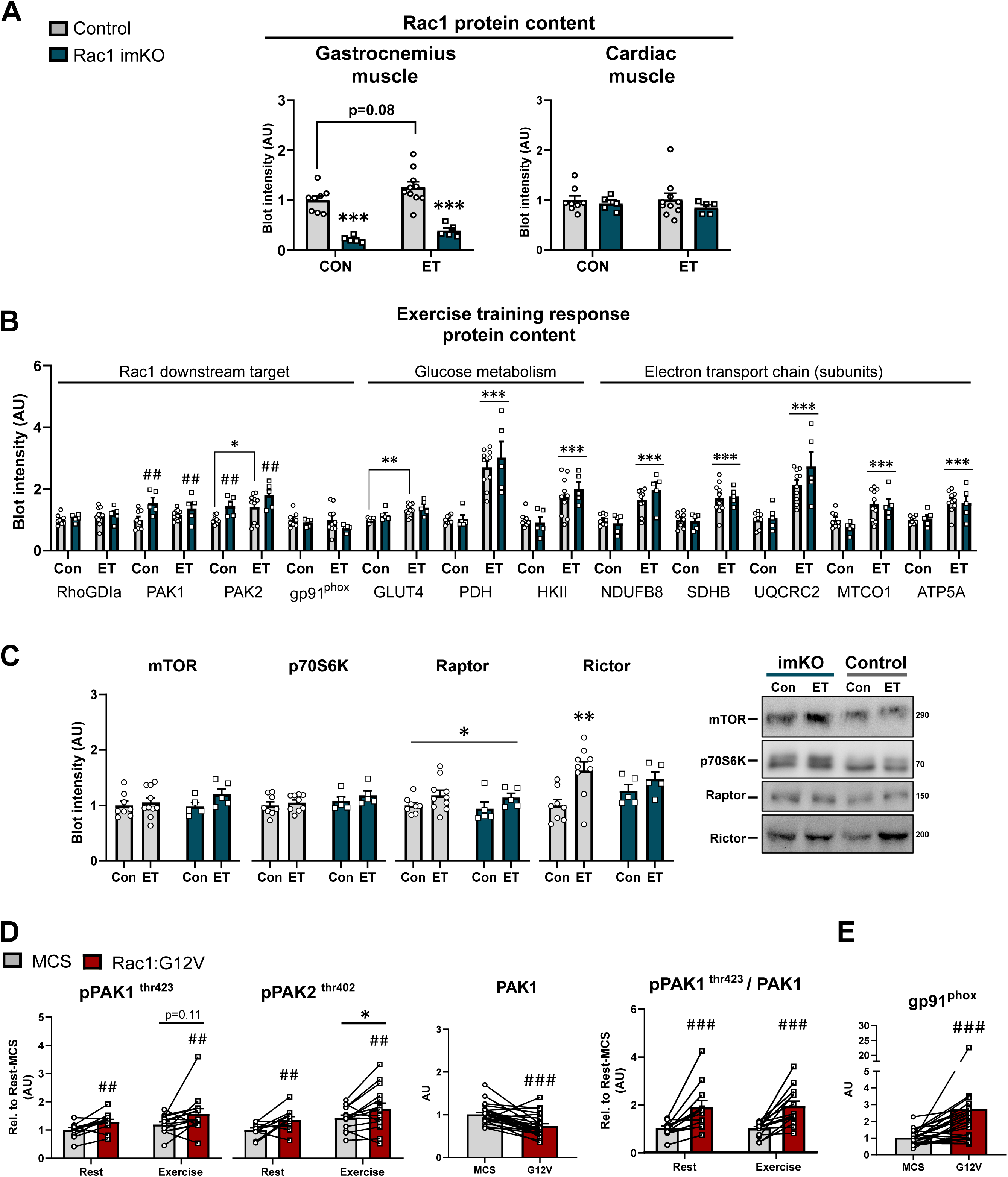
Data related to figure 6. Lack of skeletal muscle Rac1 during exercise training (ET) was investigated using inducible skeletal muscle-specific Rac1 knockout mice (Rac1 imKO) compared to control littermate mice (Con). (A) Rac1 protein content in skeletal (gastrocnemius) and cardiac muscle. (B)/(C) Effect of ET on protein content of various proteins involved in muscle metabolism and mTOR signaling in gastrocnemius muscle. (D) Effect of H-Rac1 (Rac1:G12V) on PAK1/2 signaling, and (E) gp91^phox^ protein content. For control mice: n=8-13. For Rac1 imKO mice: n= 5-13. For H-Rac1 studies: n=4-10. Significant differences of acute exercise / ET are indicated as; * = p<0.05, ** = p<0.01, *** = p<0.001. Significant differences of Rac1 imKO or H-Rac1 overexpression are indicated as; # # = p<0.01, # # # = p<0.001. Data are presented as mean + SEM incl. individual values, paired samples connected with lines when applicable.

## Methods

### Human experiments

#### Acute exercise study

These samples were acquired from a previous study. Please see ^6^ for further information.

#### One-legged kicking training study

These samples were acquired from a previous study. Please see ^38^ for further information.

### Animals

*Inducible muscle-specific Rac1 KO mice:* These mice have been described elsewhere ^15,69^. Female, age-matched, 12-16 weeks (License: 2016-15-0201-01043). Briefly, the KO was introduced by adding doxycycline to the water (1g/1L) for 3 consecutive weeks. Following 3 weeks of washout (to avoid any off-target effects of doxycycline), the mice were started on their intervention. For the exercise training study (12 weeks), all mice received two weeks of doxycycline after six weeks of voluntary wheel running to avoid potential relapse of endogenous Rac1 expression. *For NOX2 studies*: Female B10.Q control and B10.Q. p47phox mutated (*Ncf1\**) mice contain a point mutation in exon 8 generated in a previous study^70^. Age- matched non-littermate control and *Ncf1\** mice between 12 and 16 weeks of age were used for experiments (license: 2017-15-0201-01311). *For rAAV-studies:* 8 weeks old female C57BL/6 mice were commercially purchased (Taconic, Denmark). All animals were maintained on a 12:12-h light-dark cycle at 22°C ± 2°C and received a standard rodent diet (Altromin, no. 1324; Chr. Pedersen, Ringsted, Denmark) and water *ad libitum*. All experiments were approved by the Danish Animal Experimental Inspectorate.

#### Recombinant Adeno Associated Virus (rAAV) – the expression of a constitutive activated Rac1

Recombinant adeno-associated type-6 pseudotyped viral vectors (rAAV6) was created as previously described ^71^ and constructed to express a constitutively activated H-Rac1 mutant (Rac1:G12V). An intramuscular injection of the rAAV Rac1:G12V was injected in gastrocnemius (1E*10^10^) and tibialis anterior (1E*10^10^) muscle in one of the legs two weeks (acute exercise) or eight weeks (for long-term overexpression) prior to the terminal experiment. The contralateral leg was submitted to a sham treatment with an empty vector (rAAV:MCS), creating a study-design where the mice functioned as their own control (also described elsewhere ^72^).

#### Body composition

Total fat and lean body mass were measured by nuclear magnetic resonance using an EchoMRI™ (USA).

#### Treadmill exercise protocols

*<u>E</u>*<u>xercise capacity test</u> (Vmax): All mice were familiarized with the treadmill in the weeks prior to any running protocols. This consisted of 10 minutes of running at 0.16 m/s (0° incline) on three separate days. On a 15° incline, the maximal running capacity test started with a 5 min warm-up at 0.16 m/s before gradually increasing the speed by 0.02 m/s per minute until exhaustion. <u>Endurance capacity test</u>: the endurance capacity test was performed relative to the Vmax of the individual mouse. The test was performed on 10° incline. The first 5 minutes were performed at 0.10 m/s. The speed then increased to 40% of Vmax for 15 minutes and 60% of Vmax for 20 minutes. The test then continued into exhaustion gradually increasing the speed by 0.02 m/s every second minute.

#### In situ muscle contraction protocol

This protocol was developed and described elsewhere^22^. In short, mice were fasted 2 hours prior to *in situ* contraction (nine sets of contraction bouts of 1 min in duration (3 s of 10 V stimulations of pulses with a duration of 0.1 ms at a frequency of 100 Hz, repeated every 10 s), with a 30 s break between bouts). 0.2 mm acupuncture needles (TAI-CHI; B.C. Medical, Nykoebing SJ, Denmark) were inserted into the proximal and distal parts of the m. quadriceps femoris muscle. The mice were kept anaesthetized by inhalation of 2% isoflurane during the entire procedure. For the acute contraction experiment (Fig. 2A), rapamycin (1.5 mg/kg) or vehicle control (equal volume of DMSO) was injected intraperitoneally 1 h before the contraction protocol. Four hours after the contraction, the mice were anaesthetized using 2% isoflurane and given a retro-orbital injection of 21.75 mg kg−1 body weight puromycin (Calbiochem, San Diego, CA, USA) in saline to measure muscle protein synthesis^73^.

#### Glycogen measurements

Skeletal muscle glycogen was measured using the *in vitro* hexokinase method as previously described ^15^.

#### Muscle preparation and immunoblotting

Muscle tissue was pulverized in liquid nitrogen before being homogenized in a modified GSK3-buffer (10% glycerol, 1% NP-40, 20 mM sodium pyrophosphate, 150 mM NaCl, 50 mM HEPES (pH 7.5), 20 mM β-glycerophosphate, 10 mM NaF, 2 mM phenylmethylsulfonyl fluoride (PMSF), 1 mM EDTA (pH 8.0), 1 mM EGTA (pH 8.0), 2 mM Na3VO4, 10 μg/mL leupeptin, 10 μg/mL aprotinin, 3 mM benzamidine) as described previously^15^. The homogenization was performed using a Tissue-Lyser II with stainless steel grinding balls (2 x 30s at 30 Hz) (Qiagen, USA). After 30 minutes of end-over-end at 5°C, the samples were centrifuged at 9.500 *g* for 20 minutes at 4°C. The supernatant was collected discarding the remaining pellet.

Lysate protein concentration was determined using the bicinchoninic acid method. Bovine serum albumin (BSA) was used as a standard (Pierce). Immunoblotting of relevant phosphorylation-sites of proteins as well as total proteins was performed by standard immunoblotting techniques using commercially available equipment (BioRad Laboratories, USA). After the transfer of protein to polyvinylidene difluoride membranes, these were subsequently blocked for five minutes in TBS-Tween 20 containing either 2% skim milk or 3% BSA at room temperature. The membranes were incubated overnight with primary antibodies at 4°C. The primary antibodies used can be seen in Table 1. At the following day, the primary antibody was removed, and horseradish peroxidase-conjugated secondary antibody was applied and incubated for 45 minutes at room temperature. Imaging and visualization of bands were performed using Bio-Rad ChemiDocTM MP Imaging System and enhanced chemiluminescence (ECL^+^; Amersham Biosciences). For the effect of voluntary wheel running exercise training on Rac1 protein content in skeletal muscle (Figure 5B), the control mice from the Rac1 imKO exercise training cohort (12 weeks intervention, Suppl. Figure 3A), and a separate cohort (female, 6 weeks intervention) of WT mice were combined. The different cohorts are indicated with a change in color of the individual values.

**Table 1.** Overview of antibodies and dilutions.

| Antibody | Company | Number # | Dilution |
| --- | --- | --- | --- |
| Rac1 | Bd biosciences | #610651 | 1:1000 in 2% skim-milk |
| FLAG | Sigma | F1804 | 1:1000 in 2% skim-milk |
| PAK1 | Cell signaling | #2602 | 1:1000 in 2% skim-milk |
| PAK2 | Cell signaling | #2608 | 1:1000 in 2% skim-milk |
| pPAK1 thr423/<br>pPAK2 thr402 | Cell signaling | #2601 | 1:1000 in 3% BSA |
| RhoGDIa | Cell signaling | #2564 | 1:1000 in 2% skim-milk |
| Hexokinase II | Cell Signaling | #2867 | 1:1000 in 2% skim-milk |
| GLUT4 | Thermo Fisher Scientific | PA1-1065 | 1:1000 in 2% skim-milk |
| p-P38 MAPK T180/Y182 | Cell signaling | #9211 | 1:1000 in 3% BSA |
| P38 MAPK | Cell signaling | #9212 | 1:1000 in 2% skim-milk |
| p-ERK1/2 MAPK<br>T202/Y204 | Cell signaling | #9101 | 1:1000 in 2% skim-milk |
| ERK1/2 MAPK | Cell Signaling | #9102 | 1:1000 in 2% skim-milk |
| pHSP27 S82 | Cell signaling | #2406 | 1:1000 in 2% skim-milk |
| pMNK1 T197/202 | Cell signaling | #2111 | 1:1000 in 2% skim-milk |
| MNK1 | Cell Signaling | #2195 | 1:1000 in 2% skim-milk |
| pCREB S133 | Cell Signaling | #9198 | 1:1000 in 3% BSA |
| Puromycin | Sigma | MABE343 | 1:1000 in 2% skim-milk |
| p-p70S6K T389 | Cell signaling | #9205 | 1:1000 in 2% skim-milk |
| p70S6K | Cell signaling | #2708 | 1:1000 in 2% skim-milk |
| p-rS6 S235/236 | Cell Signaling | #2211 | 1:1000 in 2% skim-milk |
| rS6 | Cell signaling | #2217 | 1:1000 in 2% skim-milk |
| p-4EBP1 T37/46 | Cell signaling | #9459 | 1:1000 in 3% BSA |
| 4EBP1 | Cell Signaling | #9644 | 1:1000 in 2% skim-milk |
| gp91<sup>phox</sup> | Bd Biosciences | 611414 | 1:1000 in 2 %skim milk |
| Pyruvate dehydrogenase | Grahame Hardie, University of<br>Dundee, UK | - | 1:1000, 2% skim milk |
| Superoxide dismutase 2 | Merck Millipore | 06-984 | 1:1000 in 2% skim-milk |
| Oxphos Cocktail Rodent | Abcam | Ab110413 | 1:5000 in 3% BSA or 2%<br>skim-milk |

### RNA extraction and quantitative real-time PCR

Total RNA was extracted from 15-20 mg quadriceps using TRIzol (Qiagen, USA). One mL TRIzol was added to all samples and homogenized with stainless steel grinding balls for 2 minutes at 30 Hz using a TissueLyser II bead mill (Qiagen, USA). Then, 200 µL chloroform (Sigma, USA) was added to each sample and tubes were inverted 5 times and incubated at room temperature (RT) for 2-3 min. Following incubation, the samples were centrifuged at 8.000 RPM for 20 min at 4°C. The aqueous phase (400-500 µL) was added to 300 µL of pre-cooled isopropanol/70% ethanol (Sigma, USA), and immediately mixed via pipetting up and down 7-8 times. All samples were treated with 80 μL DNase I Incubation Mix (Qiagen, USA) and incubated at RT for 15 minutes. RNA was then further purified following the manufacturer’s protocol (Qiagen, USA). Finally, RNA was eluted by adding 30 µL RNase-free water and centrifuged at 8000 rpm for 2 min at 4°C. RNA extractions were kept on ice and their concentrations and purity were determined using a NanodropTM 2000/2000c spectrophotometer (Thermo Fisher Scientific).

Complementary (c)DNA was generated by retro-transcription reaction using the High-Capacity cDNA Reverse Transcription Kit with RNase Inhibitor (Applied Biosystems) and diluted to 10 ng/μL with nuclease-free H_2_O. mRNA content was determined using real time quantitative PCR (qPCR). Two technical replicates were analyzed per sample. qPCR was performed using the QuantStudio 6 and 7 Flex Real-Time PCR System (Applied Biosystems). The setting of the cycles used was the default for comparative Ct studies. All measurements were normalized to house- keeping mRNAs: *β-actin, sdha* or *Gapdh.* All primer sequences are listed in Table 2.

**Table 2.**
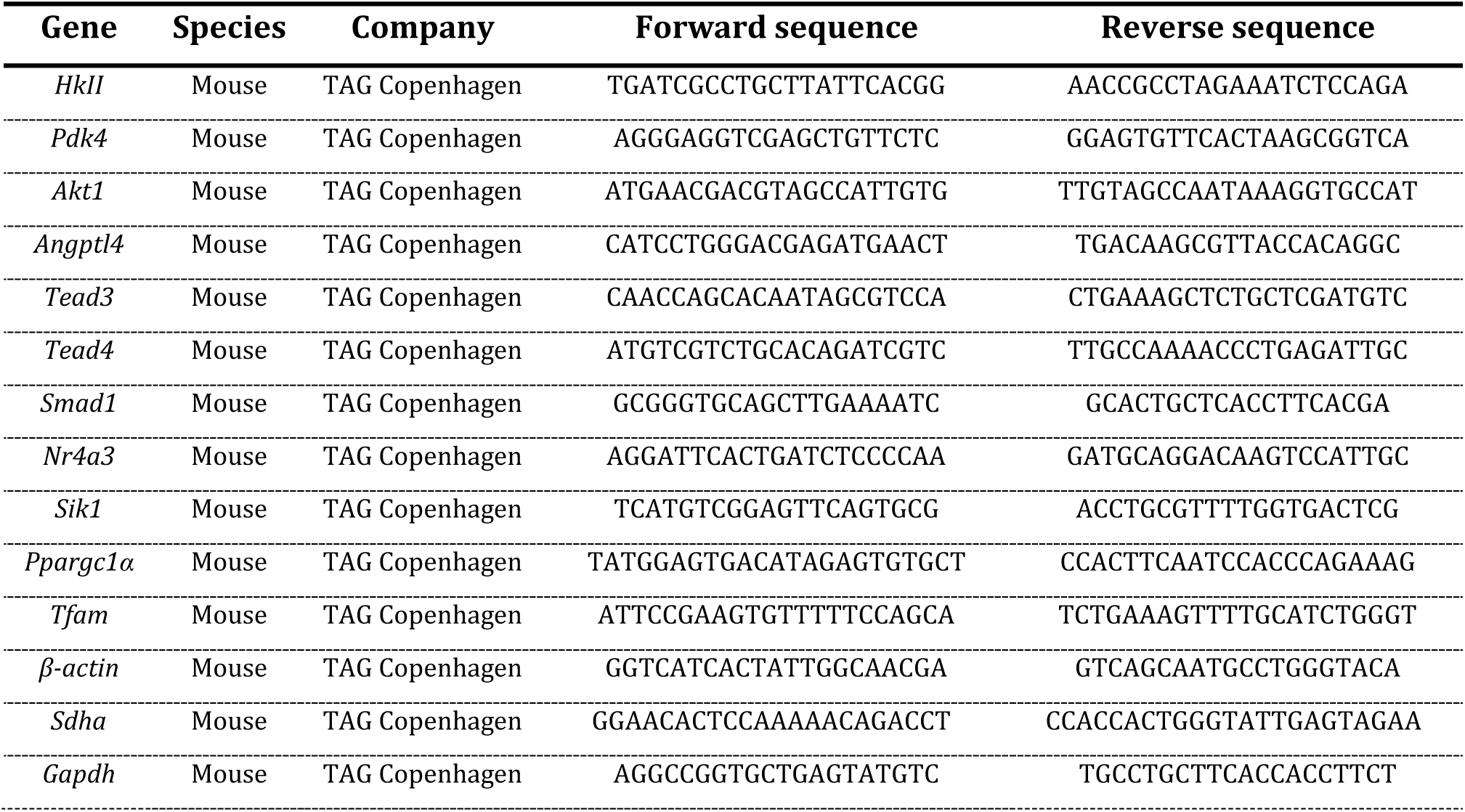
Overview of primers.

#### Total oxidant production

The staining of oxidants was performed with the commercially available 2’,7’-dichlorodihydrofluorescein diacetate (H_2_DCFDA) (#C6827, Thermo Fisher Scientific, USA). Cryo-sections (10 µm) of the embedded TA muscles were cut at -10°C and applied to microscope glasses. The samples were rinsed with PBS for five minutes before applying H_2_DCFDA (concentration). This compound is light-sensitive, and the stainings were performed with minimum light. The samples were incubated overnight at room temperature in complete darkness. The next day, any remaining H_2_DCFDA was removed before being rinsed 3x5 minutes in PBS. The samples were subsequently covered with mounting medium (H-1000, Vector Laboratories) enabling the use of laser microscopy. Confocal laser microscopy (LSM 780 confocal microscope, Zeiss) was used to visualize the production of ROS. Laser microscopy was performed blinded for interventions. The fluorescence-intensity was calculated from the average intensity of several different fibers from at least three independent pictures using ImageJ/Fiji ^74^.

#### Graphics

The graphics shown in the current study were made in ©BioRender - biorender.com (Toronto, Canada) or Inkscape©.

#### Statistical analyses

Results are shown as mean + SEM incl. individual variables when applicable. For the human and rAAV-studies, parried students t-test’s or two-way repeated measures ANOVA’s was applied when suitable. For Rac1 mKO studies, two-way ANOVA or repeated measures were applied as indicated by connecting lines. A Sidak post hoc analysis was performed if main effects during ANOVA analysis occurred. A Mann Whitney test was performed instead of a t-test, if the samples did not reach normality (Shaprio-Wilk) after log- transformation. All statistical analyses were performed using GraphPad Prism, version 9 (GraphPad Software, Inc., USA).

## References

1. Piercy, K. L. et al. The Physical Activity Guidelines for Americans. JAMA 320, 2020 (2018).

2. Hawley, J. A., Hargreaves, M., Joyner, M. J. & Zierath, J. R. Integrative Biology of Exercise. Cell 159, 738–749 (2014).

3. Neufer, P. D. et al. Understanding the Cellular and Molecular Mechanisms of Physical Activity-Induced Health Benefits. Cell Metab 22, 4–11 (2015).

4. Smith, J. A. B., Murach, K. A., Dyar, K. A. & Zierath, J. R. Exercise metabolism and adaptation in skeletal muscle. Nat Rev Mol Cell Biol (2023) doi:10.1038/s41580-023-00606-x.

5. Sylow, L. & Richter, E. A. Current advances in our understanding of exercise as medicine in metabolic disease. Curr Opin Physiol (2019) doi:10.1016/j.cophys.2019.04.008.

6. Blazev, R. et al. Phosphoproteomics of three exercise modalities identifies canonical signaling and C18ORF25 as an AMPK substrate regulating skeletal muscle function. Cell Metab 34, 1561–1577.e9 (2022).

7. Hall, A. Rho GTPases and the Actin Cytoskeleton. Science (1979) 279, 509–514 (1998).

8. Bedard, K. & Krause, K.-H. The NOX family of ROS-generating NADPH oxidases: physiology and pathophysiology. Physiol Rev 87, 245–313 (2007).

9. Abo, A. et al. Activation of the NADPH oxidase involves the small GTP-binding protein p21rac1. Nature 353, 668–70 (1991).

10. Higuchi, Y. et al. The Small GTP-binding Protein Rac1 Induces Cardiac Myocyte Hypertrophy through the Activation of Apoptosis Signal-regulating Kinase 1 and Nuclear Factor-κB. Journal of Biological Chemistry 278, 20770–20777 (2003).

11. Reil, J.-C. et al. Cardiac Rac1 overexpression in mice creates a substrate for atrial arrhythmias characterized by structural remodelling. Cardiovasc Res 87, 485–493 (2010).

12. Pracyk, J. B. et al. A requirement for the rac1 GTPase in the signal transduction pathway leading to cardiac myocyte hypertrophy. Journal of Clinical Investigation 102, 929–937 (1998).

13. Henríquez-Olguin, C. et al. Cytosolic ROS production by NADPH oxidase 2 regulates muscle glucose uptake during exercise. Nat Commun 10, 4623 (2019).

14. Henríquez-Olguín, C. et al. The Emerging Roles of Nicotinamide Adenine Dinucleotide Phosphate Oxidase 2 in Skeletal Muscle Redox Signaling and Metabolism. Antioxid Redox Signal 31, 1371–1410 (2019).

15. Sylow, L. et al. Rac1 governs exercise-stimulated glucose uptake in skeletal muscle through regulation of GLUT4 translocation in mice. J Physiol 17, 4997–5008 (2016).

16. Sylow, L. et al. Rac1 is a novel regulator of contraction-stimulated glucose uptake in skeletal muscle. Diabetes 62, 1139–1151 (2013).

17. Manser, E., Leung, T., Salihuddin, H., Zhao, Z. & Lim, L. A brain serine/threonine protein kinase activated by Cdc42 and Rac1. Nature 367, 40–46 (1994).

18. Manser, E. et al. Molecular cloning of a new member of the p21-Cdc42/Rac-activated kinase (PAK) family. Journal of Biological Chemistry 270, 25070–25078 (1995).

19. Tsakiridis, T., Taha, C., Grinsteinl, S. & Klip, A. Insulin activates a p21-activated kinase in muscle cells via phosphatidylinositol 3-kinase. Journal of Biological Chemistry 271, 19664– 19667 (1996).

20. Sylow, L. et al. Akt and Rac1 signaling are jointly required for insulin-stimulated glucose uptake in skeletal muscle and downregulated in insulin resistance. Cell Signal 26, 323–331 (2014).

21. Sylow, L. et al. Rac1 signaling is required for insulin-stimulated glucose uptake and is dysregulated in insulin-resistant murine and human skeletal muscle. Diabetes 62, 1865–1875 (2013).

22. Knudsen, J. R. et al. Contraction-regulated mTORC1 and protein synthesis: Influence of AMPK and glycogen. J Physiol 598, 2637–2649 (2020).

23. Steinert, N. D. et al. Mapping of the contraction-induced phosphoproteome identifies TRIM28 as a significant regulator of skeletal muscle size and function. Cell Rep 34, 108796 (2021).

24. Ryder, J. W. et al. Effect of Contraction on Mitogen-activated Protein Kinase Signal Transduction in Skeletal Muscle. Journal of Biological Chemistry 275, 1457–1462 (2000).

25. Zheng, C. et al. MAPK-activated Protein Kinase-2 (MK2)-mediated Formation and Phosphorylation-regulated Dissociation of the Signal Complex Consisting of p38, MK2, Akt, and Hsp27. Journal of Biological Chemistry 281, 37215–37226 (2006).

26. Fukunaga, R. MNK1, a new MAP kinase-activated protein kinase, isolated by a novel expression screening method for identifying protein kinase substrates. EMBO J 16, 1921– 1933 (1997).

27. Waskiewicz, A. J. Mitogen-activated protein kinases activate the serine/threonine kinases Mnk1 and Mnk2. EMBO J 16, 1909–1920 (1997).

28. Xing, J., Ginty, D. D. & Greenberg, M. E. Coupling of the RAS-MAPK Pathway to Gene Activation by RSK2, a Growth Factor-Regulated CREB Kinase. Science (1979) 273, 959– 963 (1996).

29. Knudsen, J. R. et al. Exercise increases phosphorylation of the putative mTORC2 activity readout NDRG1 in human skeletal muscle. American Journal of Physiology-Endocrinology and Metabolism 322, E63–E73 (2022).

30. Kumar, V., Atherton, P., Smith, K. & Rennie, M. J. Regulation of Protein Metabolism in Exercise and Recovery Human muscle protein synthesis and breakdown during and after exercise. J Appl Physiol 106, 2026–2039 (2009).

31. Ogasawara, R., Knudsen, J. R., Li, J., Ato, S. & Jensen, T. E. Rapamycin and mTORC2 inhibition synergistically reduce contraction-stimulated muscle protein synthesis. J Physiol JP280528 (2020) doi:10.1113/JP280528.

32. Ogasawara, R. & Suginohara, T. Rapamycin-insensitive mechanistic target of rapamycin regulates basal and resistance exercise-induced muscle protein synthesis. The FASEB Journal 32, 5824–5834 (2018).

33. Roberts, M. D. et al. Mechanisms of mechanical overload-induced skeletal muscle hypertrophy: current understanding and future directions. Physiol Rev (2023) doi:10.1152/physrev.00039.2022.

34. Hingst, J. R. et al. Exercise-induced molecular mechanisms promoting glycogen supercompensation in human skeletal muscle. Mol Metab 16, 24–34 (2018).

35. Satoh, M. et al. Requirement of Rac1 in the development of cardiac hypertrophy. Proc Natl Acad Sci U S A 103, 7432–7 (2006).

36. Gelderman, K. A., Hultqvist, M., Holmberg, J., Olofsson, P. & Holmdahl, R. T cell surface redox levels determine T cell reactivity and arthritis susceptibility. Proceedings of the National Academy of Sciences 103, 12831–12836 (2006).

37. Sareila, O., Jaakkola, N., Olofsson, P., Kelkka, T. & Holmdahl, R. Identification of a region in p47phox/NCF1 crucial for phagocytic NADPH oxidase (NOX2) activation. J Leukoc Biol 93, 427–435 (2012).

38. Frøsig, C. et al. Effects of Endurance Exercise Training on Insulin Signaling in Human Skeletal Muscle. Diabetes 56, 2093–2102 (2007).

39. Kleinert, M. et al. Quantitative proteomic characterization of cellular pathways associated with altered insulin sensitivity in skeletal muscle following high-fat diet feeding and exercise training. Sci Rep 8, 10723 (2018).

40. Manzanares, G., Brito-da-Silva, G. & Gandra, P. G. Voluntary wheel running: patterns and physiological effects in mice. Braz J Med Biol Res 52, e7830 (2018).

41. Raun, S. H. et al. Housing temperature influences exercise training adaptations in mice. Nat Commun 11, 1560 (2020).

42. McKie, G. L. et al. Housing temperature affects the acute and chronic metabolic adaptations to exercise in mice. J Physiol 597, 4581–4600 (2019).

43. Møller, L. L. V. et al. The Rho guanine dissociation inhibitor α inhibits skeletal muscle Rac1 activity and insulin action. Proceedings of the National Academy of Sciences 120, (2023).

44. Egan, B. & Sharples, A. P. Molecular responses to acute exercise and their relevance for adaptations in skeletal muscle to exercise training. Physiol Rev 103, 2057–2170 (2023).

45. Furrer, R., Hawley, J. A. & Handschin, C. The molecular athlete: exercise physiology from mechanisms to medals. Physiol Rev 103, 1693–1787 (2023).

46. Watt, K. I. et al. The Hippo pathway effector YAP is a critical regulator of skeletal muscle fibre size. Nat Commun 6, 6048 (2015).

47. Bid, H. K., Roberts, R. D., Manchanda, P. K. & Houghton, P. J. RAC1: An Emerging Therapeutic Option for Targeting Cancer Angiogenesis and Metastasis. Mol Cancer Ther 12, 1925–1934 (2013).

48. Prior, I. a, Lewis, P. D. & Mattos, C. A comprehensive survey of Ras mutations in cancer. Cancer Res 72, 2457–2467 (2012).

49. Kawazu, M. et al. Transforming mutations of RAC guanosine triphosphatases in human cancers. Proceedings of the National Academy of Sciences 110, 3029–3034 (2013).

50. Sylow, L., Møller, L. L. V., Kleinert, M., Richter, E. A. & Jensen, T. E. Stretch-stimulated glucose transport in skeletal muscle is regulated by Rac1. J Physiol 593, 645–656 (2015).

51. Saxton, R. A. & Sabatini, D. M. mTOR Signaling in Growth, Metabolism, and Disease. Cell 168, 960–976 (2017).

52. Saci, A., Cantley, L. C. & Carpenter, C. L. Rac1 Regulates the Activity of mTORC1 and mTORC2 and Controls Cellular Size. Mol Cell 42, 50–61 (2011).

53. Akimoto, T. et al. Exercise Stimulates Pgc-1α Transcription in Skeletal Muscle through Activation of the p38 MAPK Pathway. Journal of Biological Chemistry 280, 19587–19593 (2005).

54. Pogozelski, A. R. et al. p38γ Mitogen-Activated Protein Kinase Is a Key Regulator in Skeletal Muscle Metabolic Adaptation in Mice. PLoS One 4, e7934 (2009).

55. Handschin, C., Rhee, J., Lin, J., Tarr, P. T. & Spiegelman, B. M. An autoregulatory loop controls peroxisome proliferator-activated receptor γ coactivator 1α expression in muscle. Proceedings of the National Academy of Sciences 100, 7111–7116 (2003).

56. Kammoun, M. et al. The Invalidation of HspB1 Gene in Mouse Alters the Ultrastructural Phenotype of Muscles. PLoS One 11, e0158644 (2016).

57. Bruno, N. E. et al. Creb coactivators direct anabolic responses and enhance performance of skeletal muscle. EMBO J 33, 1027–1043 (2014).

58. Bramham, C. R., Jensen, K. B. & Proud, C. G. Tuning Specific Translation in Cancer Metastasis and Synaptic Memory: Control at the MNK–eIF4E Axis. Trends Biochem Sci 41, 847–858 (2016).

59. Ueda, T., Watanabe-Fukunaga, R., Fukuyama, H., Nagata, S. & Fukunaga, R. Mnk2 and Mnk1 Are Essential for Constitutive and Inducible Phosphorylation of Eukaryotic Initiation Factor 4E but Not for Cell Growth or Development. Mol Cell Biol 24, 6539–6549 (2004).

60. Sandeman, L. Y. et al. Disabling MNK protein kinases promotes oxidative metabolism and protects against diet-induced obesity. Mol Metab 42, 101054 (2020).

61. Cerquone Perpetuini, A., et al. Group I Paks support muscle regeneration and counteract cancer-associated muscle atrophy. J Cachexia Sarcopenia Muscle 9, 727–746 (2018).

62. Joseph, G. A. et al. Group I Paks Promote Skeletal Myoblast Differentiation In Vivo and In Vitro. Mol Cell Biol 37, (2017).

63. Joseph, G. A. et al. Late-onset megaconial myopathy in mice lacking group I Paks. Skelet Muscle 9, 5 (2019).

64. Henriquez-Olguin, C. et al. NOX2 deficiency exacerbates diet-induced obesity and impairs molecular training adaptations in skeletal muscle. Redox Biol 65, 102842 (2023).

65. D’Hulst, G., Masschelein, E. & De Bock, K. Resistance exercise enhances long-term mTORC1 sensitivity to leucine. Mol Metab 66, 101615 (2022).

66. Davis, R. T. et al. Knockout of p21-activated kinase-1 attenuates exercise-induced cardiac remodelling through altered calcineurin signalling. Cardiovasc Res 108, 335–347 (2015).

67. Henríquez-Olguín, C. et al. Adaptations to high-intensity interval training in skeletal muscle require NADPH oxidase 2. Redox Biol 24, 101188 (2019).

68. Margaritelis, N. V. et al. Adaptations to endurance training depend on exercise-induced oxidative stress: exploiting redox interindividual variability. Acta Physiologica 222, e12898 (2018).

69. Raun, S. H. et al. Rac1 muscle knockout exacerbates the detrimental effect of high-fat diet on insulin-stimulated muscle glucose uptake independently of Akt. J Physiol 596, 2283–2299 (2018).

70. Hultqvist, M. et al. Enhanced autoimmunity, arthritis, and encephalomyelitis in mice with a reduced oxidative burst due to a mutation in the *Ncf1* gene. Proceedings of the National Academy of Sciences 101, 12646–12651 (2004).

71. Blankinship, M. J. et al. Efficient transduction of skeletal muscle using vectors based on adeno-associated virus serotype 6. Molecular Therapy 10, 671–678 (2004).

72. Blankinship, M. J. et al. Efficient transduction of skeletal muscle using vectors based on adeno-associated virus serotype 6. Molecular Therapy 10, 671–678 (2004).

73. Goodman, C. A. et al. Novel insights into the regulation of skeletal muscle protein synthesis as revealed by a new nonradioactive *in vivo* technique. The FASEB Journal 25, 1028–1039 (2011).

74. Schindelin, J., et al. Fiji: an open-source platform for biological-image analysis. Nat Methods 9, 676–82 (2012).

